# DeepImmuno: Deep learning-empowered prediction and generation of immunogenic peptides for T cell immunity

**DOI:** 10.1101/2020.12.24.424262

**Authors:** Guangyuan Li, Balaji Iyer, V. B. Surya Prasath, Yizhao Ni, Nathan Salomonis

**Author notes:** **Corresponding Author**: Guangyuan Li.

## Abstract

T-cells play an essential role in the adaptive immune system by seeking out, binding and destroying foreign antigens presented on the cell surface of diseased cells. An improved understanding of T-cell immunity will greatly aid in the development of new cancer immunotherapies and vaccines for life threatening pathogens. Central to the design of such targeted therapies are computational methods to predict non-native epitopes to elicit a T cell response, however, we currently lack accurate immunogenicity inference methods. Another challenge is the ability to accurately simulate immunogenic peptides for specific human leukocyte antigen (HLA) alleles, for both synthetic biological applications and to augment real training datasets. Here, we proposed a beta-binomial distribution approach to derive epitope immunogenic potential from sequence alone. We conducted systematic benchmarking of five traditional machine learning (ElasticNet, KNN, SVM, Random Forest, AdaBoost) and three deep learning models (CNN, ResNet, GNN) using three independent prior validated immunogenic peptide collections (dengue virus, cancer neoantigen and SARS-Cov-2). We chose the CNN model as the best prediction model based on its adaptivity for small and large datasets, and performance relative to existing methods. In addition to outperforming two highly used immunogenicity prediction algorithms, DeepHLApan and IEDB, DeepImmuno-CNN further correctly predicts which residues are most important for T cell antigen recognition. Our independent generative adversarial network (GAN) approach, DeepImmuno-GAN, was further able to accurately simulate immunogenic peptides with physiochemical properties and immunogenicity predictions similar to that of real antigens. We provide DeepImmuno-CNN as source code and an easy-to-use web interface.

**Data Availability:** DeepImmuno Python3 code is available at https://github.com/frankligy/DeepImmuno. The DeepImmuno web portal is available from https://deepimmuno.herokuapp.com. The data in this article is available in GitHub and supplementary materials.

## INTRODUCTION

Immunotherapy has emerged as a promising strategy to combat cancer by “reprogramming” a patient’s own immune system. Effective targeted immunotherapies require accurately predicting which cancer-specific neo-epitopes are most likely to elicit an immune response. Similar strategies are currently being designed to target antigens commonly produced by serious pathogens, such as the SARS-Cov-2 (COVID-19) virus [1]. Human leukocyte antigens (HLAs) are a polymorphic class of proteins on the cell surface of T cells that recognize foreign antigens presented by another cell. The process of antigen recognition is the cornerstone of the adaptive immune system. HLA proteins are encoded by the Major Histocompatibility Complex (MHC) genes in humans. Predicting the immunogenicity of MHC-I bound epitopes is crucial for understanding the molecular rules governing T cell directed adaptive immunity and creating precision cancer or pathogen targeting vaccines. Cellular antigen recognition is governed by a series of carefully orchestrated molecular interactions between cell-surface-presented antigen and T cells of the immune system. MHC-I proteins are responsible for presentation of short epitopes on the cell surface and mediating interactions with CD8+ T cell receptors (TCR). An immunogenic peptide is capable of binding with a cognate MHC molecule, resulting in the exposure of its non-self portion. The exposure of “foreign” signals trigger immunoreceptor tyrosine-based activation motifs (ITAMs) on the T cell to be phosphorated and activate an immune response [2]. The process ultimately results in targeted cell death of the antigen expression cell by CD8 T cell. Hence, the identification of immunogenic epitopes that can trigger T cell responses is central to developing new cancer immunotherapies and vaccines. Because thousands of potential disease-associated antigens can be presented in innate or foreign cells [3], it is necessary to prioritize which candidates are most likely to induce T cell response prior to experimental validation.

To reduce the number of epitopes to be chosen, *in silico* methods have been developed to predict antigen immunogenicity. POPI [4] was developed as the first automated computational immunogenicity prediction tool. POPI used a selected subset of physicochemical features identified by a bi-objective algorithm for support vector machine (SVM) based classification. An updated version POPISK [5] further considers MHC binding properties to improve its prediction ability. PAAQD [6] was later developed to consider amino acid pairwise contact potential and quantum topological molecular similarity (QTMS) for feature selection. Subsequently, a machine learning-based immunogenicity predictor NeoPepsee [7] was developed that integrated 14 independent features to infer peptide immunogenicity. These initial methods paved the way for more advanced algorithms, however, the applicability of such methods have historically been challenging due to small training datasets and limited consideration of HLA alleles. A significant advance in the field came with the introduction of the immune epitope database (IEDB) and associated predictive immunogenicity tools [8]. This invaluable resource continues to systematically characterize the biochemical properties of over 30,000 MHCI-bound immunogenic epitopes. IEDB further includes a suite of algorithms to predict binding affinity and immunogenicity, including a position-weighted calculated schema by considering kullback-leibler (KL) divergence and amino acid preference (default method). More recently, algorithms with improved reported accuracy have been described, including a Random Forest based approach called INeo-Epp [9] which uses a customized immunogenic score and the recurrent neural network-based deep learning approach DeepHLApan [10]. While promising, a potential limitation of these these approaches is that the prediction of immunogenic epitopes is treated as a binary classification problem using predefined hard cutoffs, in which each peptide-MHC pair will be considered immunogenic or non-immunogenic, even though the immunogenicity of a certain peptide-MHC will vary substantially depending on the subject’s immune profile and TCR repertoire [2]. Further, while DeepHLApan [10] applies a wellrationaled deep learning approach, its encoding of amino-acid sequence does not incorporate physicochemical or other amino-acid parameters (one-hot encoding). As a result, the outputs from these methods might not fully reflect the ability of the peptide-MHC to trigger a T cell response.

A secondary, but important challenge in the field of immunogenicity prediction, is to learn the rules that govern which peptides are immunogenic and why. Understanding these rules, could be used to develop improved prediction models or produce large synthetic datasets for training more accurate predictive models. Deep generative models [11] are a newly-emerging area in artificial intelligence (AI) that can be applied to diverse research problems. In effect, such models allow for the creation of accurate synthetic models from limited existing training data. Such methods take random noise to create new datasets that reflect the original training data but that contain unique informative features. Generative adversarial networks (GANs) are widely used in computer vision [12] and synthetic biology [13] to generate new images or sequences of interest (i.e. antimicrobial peptides), but have not previously used to produce synthetic models of immunogenic peptides.

To overcome the aforementioned limitations, we propose a new convolutional neural network (CNN) [14] approach called DeepImmuno-CNN. During the training, a beta-binomial probabilistic model is fitted to the training dataset to derive a continuous immunogenic score. This score differentially weights each peptide-MHC complex in the model based on its associated experimental evidence (high-confidence or low-confidence), to further produce a more reliable variable immunogenic score for each peptide in the test dataset. Each amino acid sequence is additionally encoded using a reduced principal component analysis (PCA) feature space of 566 well-curated amino acid physicochemical features from the AAindex1 database [15] to overcome sparsity issues related to one-hot encoding [16]. Diverse machine learning and deep learning approaches exist, which have potential strengths and weaknesses for this problem (e.g., performance, accuracy, flexibility to dataset size). To ensure the rigor of this approach, we performed a systematic comparison of five traditional machine learning algorithms (ElasticNet, K-Nearest Neighbors (KNN), SVM, Random Forest, AdaBoost) and three deeplearning models (CNN, Graph Neural Network (GNN), Residual Net (ResNet)). This benchmarking further supports the use of a CNN for this problem. In addition, an evaluation of different encoding schemas, confirms that our AAindex1 PCA encoding strategy provides excellent performance relative to alternative methods. When benchmarked against two state-of-the-art workflows for immunogenicity prediction (DeepHLApan and IEDB), DeepImmuno-CNN was able to significantly increase both precision and recall for different HLA genotypes using diverse real-world test datasets (IEDB, TESLA and COVID-19). To further explore the dependent epitope features for immunogenicity prediction, we developed a GAN model [13][17] which mimics the salient features of validated immunogenic peptides. These data support the hypothesis that immunogenic peptides are learnable as a possible future source for high quality synthetic training data.

Hence, this work represents multiple important advances and insights into the field of immunogenicity prediction, including: 1) comprehensive benchmarking of existing and new methods, 2) improved quantitative prediction models, 3) applicability for neoantigen and infectious peptides, 4) crucial determinants for T cell responses and 5) an accurate approach for synthetic modeling.

## METHODS

### Datasets

Multiple training and test datasets were analyzed in this study using previously published experimentally tested datasets. For initial training and validation, we analyzed >9,000 experimentally evaluated immunogenicity assay predictions from the Immune Epitope Database, IEDB database (August 13th, 2020). For our evaluation, we restricted the dataset to peptides with metadata that matched to the following keywords: (1) linear epitope, (2) T cell assay, (3) MHC class I, (4) human, and (5) disease. To restrict the dataset to informative predictions, we developed a rigorous data cleaning strategy. First, data instances without explicit 4-digit MHC alleles were discarded. Second, all redundant peptide-MHC allele instances were discarded (the same peptide with different HLA alleles were considered different instances). Third, all negative epitopes, without explicit experimental information (number of subjects tested, number of subjects responded) or with less than four tested subjects were removed (likely not informative at a human population-level). Fourth, peptides of length 9 and 10 were retained for the training process. 9-mer and 10-mer peptides cover 97.5% of all data instances and are also the dominant length for MHCl-bound peptides [18]. Finally, we separated out 408 dengue virus positive instances from Weiskopf et al [19] for the purpose of internal validation of different prediction methods. Specifically, 9,056 data instances were retained in the final dataset, among which 4,143 were positive reactive instances and the remaining 4,913 were negative. We used ten-fold cross validation for internal benchmark analysis to avoid over-fitting. That is, we split the datasets into 10 rotating subsets – nine for training and one for validation in each run. At the end of cross validation, the scores for each evaluation metric were averaged over the ten testing subsets as the model’s performance. We selected two independent test datasets for further evaluation: 1) 637 experimentally tested tumor specific neoantigens from the Tumor Neoantigen Selection Alliance (TESLA) [20], and 2) 100 SARS-Cov-2 peptides [1,20] tested for their immunogenicity in convalescent and unexposed subjects, respectively.

### Encoding Strategy

To represent each HLA allele and encoded peptide sequences in a numerical matrix as the input for each evaluated machine learning and deep learning algorithms, we developed and tested different encoding strategies. We used HLA paratopes (HLA-antigen interacting residues) as a proxy of different HLA alleles as these sequences contain the most salient information to describe peptide-HLA spatial interactions. The AAindex encoding strategy was designed to account for amino acid comprehensive physicochemical properties.

*AAindex:* We retrieved 566 amino acid associated physicochemical properties from the AAindex1 database [15]. Among the 566 properties, 13 indices were discarded due to missing values for certain amino acids (**Supplementary Table 1**). We introduced a placeholder amino acid “-” for padding the gaps of HLA paratope sequences and 9-mer peptides (see below). The corresponding AAindex values were set as the average of all other 20 canonical amino acids. This method adds the total amino acid number to 21. The resulting 21 x 553 numeric matrix was normalized using RobustScaler [21] via the following operation:

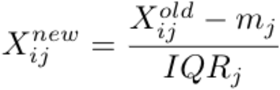

Where **X** is the numeric matrix, **m** is the median per each feature column and **IQR** is the interquartile (Q3-Q1) per each feature column. The normalized feature matrix undergoes a principal component analysis (PCA) to remove noisy features such that it only retains relevant components. We chose 12 principal components which explain 95% total variance. This step leads to a 21 x 12 numerical matrix (hereafter AAindex matrix). For peptides, we adopted an encoding schema similar to that of O’Donnell et al [22] to pad shorter peptides (9-mer) to a longer sequence (10-mer) such that first five residues and the last four residues were joined by a placeholder “-”, since the two termini are often involved in binding interactions [23,24]. For MHC molecules, we encoded each MHC allele based on its parotopes sequence, which is the set of discontinuous residues sterically interacting with peptides. This paratope information was evidenced and analyzed from crystal structure and was retrieved from IMGT-3D-Structure database [25] (http://www.imgt.org/3Dstructure-DB/). For MHC alleles which did not have a solved peptide-MHC structure, their paratope information was determined by their neighbors. Specifically, the paratopes of allele HLA-A*2403 were determined by its nearest neighbor HLA-A*2402, which is a more frequent allele whose paratope sequence is available. We then performed two rounds of multiple sequence alignment using clustal-omega [26]. The first iteration was used for generating a consensus sequence for a single HLA allele from all its solved crystal structure, while the second round was for all paratope sequences with same length, gaps were filled with the placeholder [26]. A schematic example is shown in **Supplementary Figure 1**.

**Supplementary Figure 1.**
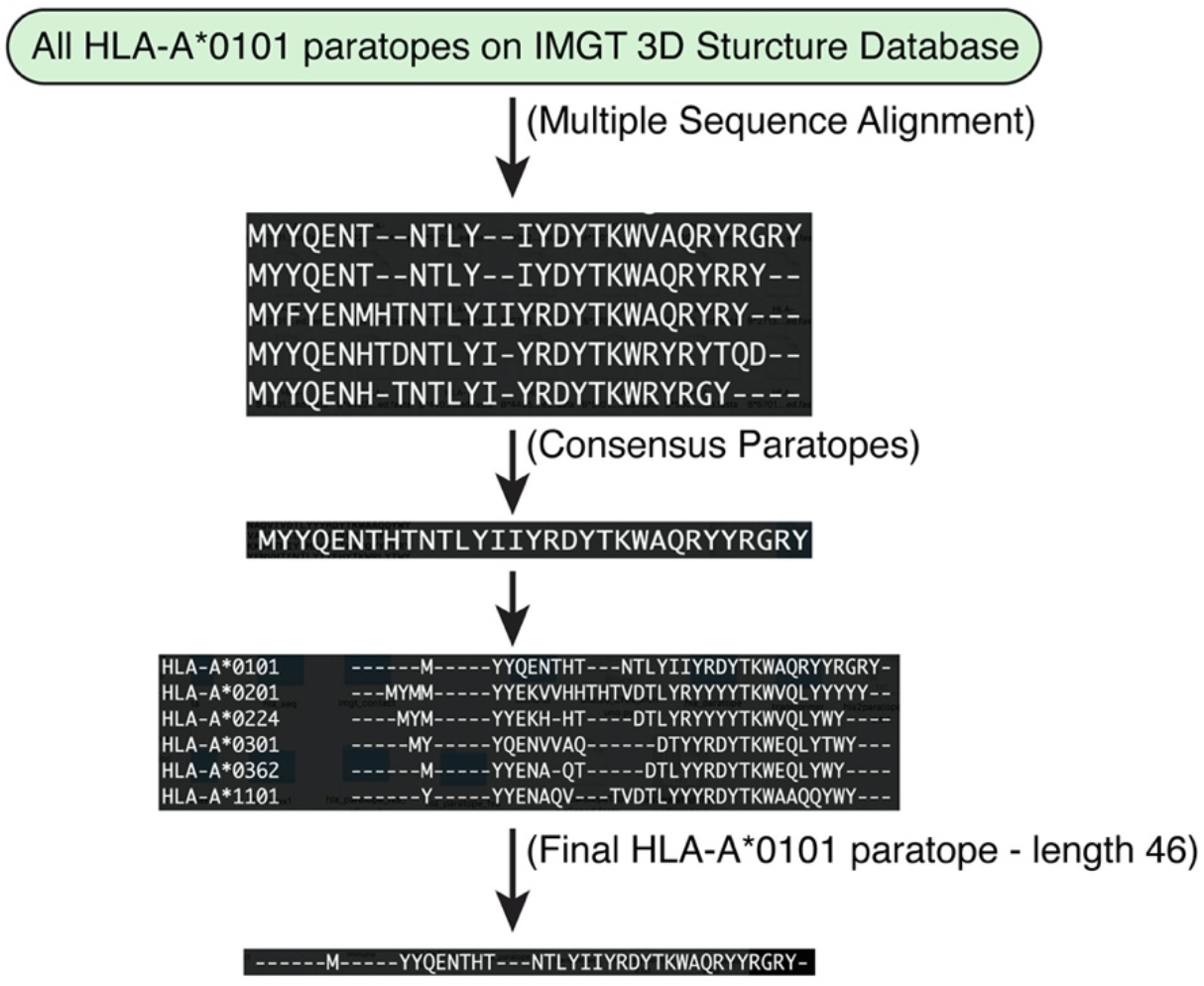
Workflow to generate HLA paratope sequences. To predict the antigen-binding residues of each HLA allele or paratope, we constructed consensus paratope sequences for each known human HLA allele. Shown here are five example sequences from five independent solved crystal structures of the HLA allele HLA-A*0101. A consensus paratope sequence was determined by computing the most frequent residue in each position. Another round of multiple sequence alignment generates fixed-length HLA paratopes for all HLA alleles. The token “-” is introduced to represent nicks and gaps.

### Beta Binomial immunogenic model

Three columns of information from the IEDB database were used in the creation of the beta-binomial model, namely the immunogenic class (**x**), result claimed by submitter (positive, positive-high, positive-intermediate, positive-low, negative), number of subjects tested (**s**) and number of subjects responded (**s-f**). We derived a prior beta distribution based on the immunogenic class (**x**):

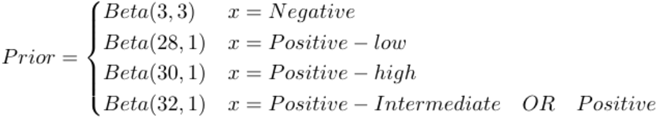

For a given epitope (data instance), assuming that we observe **s** successful T cell responses and **f** fails, then the posterior distribution of this epitope’s immunogenic potential follow a new beta distribution:

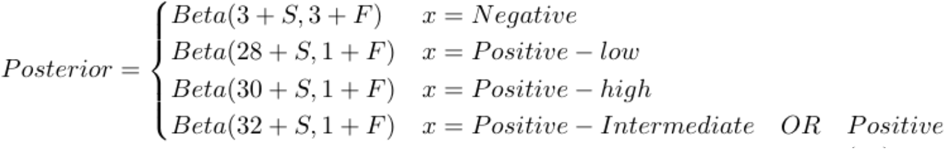

We then performed 50 bootstrapped iterations from the derived posterior distribution and used the average as the final immunogenic potential of a certain peptide-MHC complex.

### Prediction models

We first adopted and rigorously compared the performance of five machine learning algorithms (ElasticNet, KNN, SVM, Random Forest and AdaBoost), after optimizing parameters for each method as follows. ElasticNet regression was first cross-validated to determine the best hyperparameters (alpha=0.01, l1_ratio=0.51), where alpha controlled the regularization strength and l1_ratio determined the percentage of the L1-norm penalty (lasso regression) and the L2-norm penalty (ridge regression). KNN regressor was cross-validated to determine the best hyperparameters (n_neighbors=23), n_neighbors control the neighbor information used for inferring query point’s properties. SVM linear regressor was cross-validated to determine the best hyperparameter (C=0.01), C is the reciprocal of regularization strength which is inversely proportional to how many mistakes are allowed in the model. Random Forest was crossvalidated to determine the best hyperparameter (n_estimators=200, min_sample_leaf=1), n_estimators control the number of decision trees in the model and min_sample_leaf control the minimum amount of samples to be a leaf node. The aforementioned cross-validations were all 10-fold and rooted mean square error (RMSE) was used as default evaluation criteria if not specifically mentioned otherwise. The same hyperparameters were adopted for the adaptive boost (AdaBoost) model, similar to Random Forest, since they are both tree-based ensemble methods.

We further implemented and optimized three deep learning architectures:

*CNN:* The pictorial architecture is shown in **Figure 1C**. Peptide and MHC were processed by two consecutive convolutional layers, followed by two dense layers to consider the interactions between peptide and MHC. The basic convolution operation is mathematically represented as:

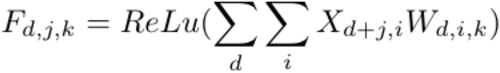

**Figure 1.**
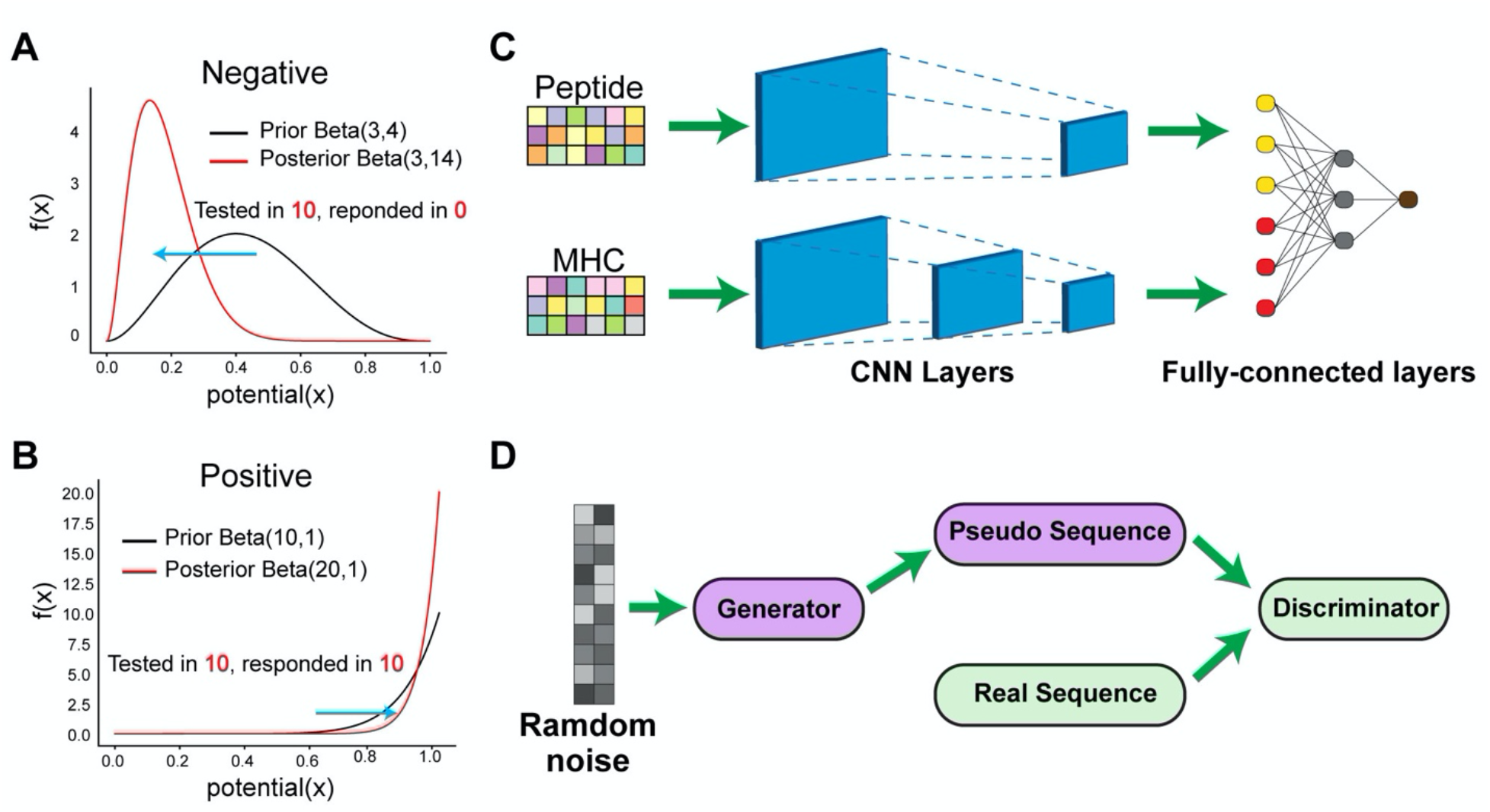
The DeepImmuno model. (A-B) In DeepImmuno, to assess the probability that a given antigen is immunogenic, variable peptide immunogenic potential is computed by sampling from a posterior beta distribution of well-defined true positive and true negative immunogenic antigens to produce a continuous immunogenic score. The posterior distribution is derived using a subset of T-cell immunogenic assay results from the Immune Epitope DataBase (binomial) and a prior beta distribution of either (A) negative or (B) positive assay results. (C) The DeepImmuno-CNN architecture is shown to predict interactions between each peptide and MHC allele. In this model, each peptide/MHC pair is subjected to two consecutive convolutional layers, followed by two fully-connected dense layers to output a predictive value for each pair. (D) The Deeplmmuno-GAN architecture is depicted for simulating immunogenic peptide sequences using only random sequences as an input. The GAN model is composed of a generator and a discriminator. This learning generator produces pseudo-sequences in an attempt to artificially convince the discriminator the immunogenic sequences are real, while the discriminator uses real peptides sequences along with generated pseudo-sequences to distinguish the difference.

Where **F** is the resultant feature map, **X** is the input numerical matrix and **W** is the kernel. Lower case **d** denotes the row index and **i** denotes the column index of original matrix and **k** denotes the index of kernels in the convolutional layers. The **ReLu** function was used as an activation function. When training the model, we set the batch_size = 128. Two early stopping strategies were adopted: 1) monitor the training_loss with patience = 2; training will immediately stop if training loss increases and 2) monitor the valiation_loss with patience=15; training will stop when we did not observe validation loss decrease in 15 epochs.

*ResNet:* An overview is shown in **Supplement Figure 2A**. Peptide and MHC undergo three consecutive residue blocks, each residual block containing three CNN layers followed by a maxpool layer. Two dense layers were used at the end for prediction. Each residual block [27] contains skip connection which feed the input back to the output to avoid gradient vanishing as determined by:

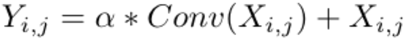

Where **Y** is the output matrix of a single residual block, α determines the fraction of convolutional output we want to keep, **X** is the original input matrix.

*GNN:* An overview is shown in **Supplementary Figure 2B**. Each peptide-MHC complex was represented by an acyclic undirected graph. Two types of edges were specified, ones were intra-edges denoting the interactions between/within-peptide and within-MHC interactions, others were inter-edges denoting the interactions between peptide and MHC. To emphasize the peptide-MHC interactions, we assigned a weight = 2 on inter-edges and weight = 1 on intra-edges. Two graph convolutional layers [28] were built upon the constructed graph objects, followed by a mean readout layer [29] to summarize node embedding at the graph level. The learned graph level features are fed into two dense layers for predictions. The core graph convolution operation can be mathematically described as:

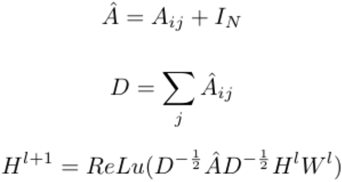

Where **A_ij_** is the adjacency matrix of graph objects, **i-**th row and **j-**th column represent the **i**th node and its **j-**th associated feature and **I**_N_ is the self-loop which is a diagonal matrix. The degree matrix **D** is the sum of adjacency matrix over the columns. **H** is the graph representation, which corresponds to a **N** x **M** matrix where **N** is the number of nodes and **M** is the number of features associated with each node. Lower case **i** denotes the layer of graph representation and **W** is the trainable weight matrix that governs the learning process.

**Supplementary Figure 2.**
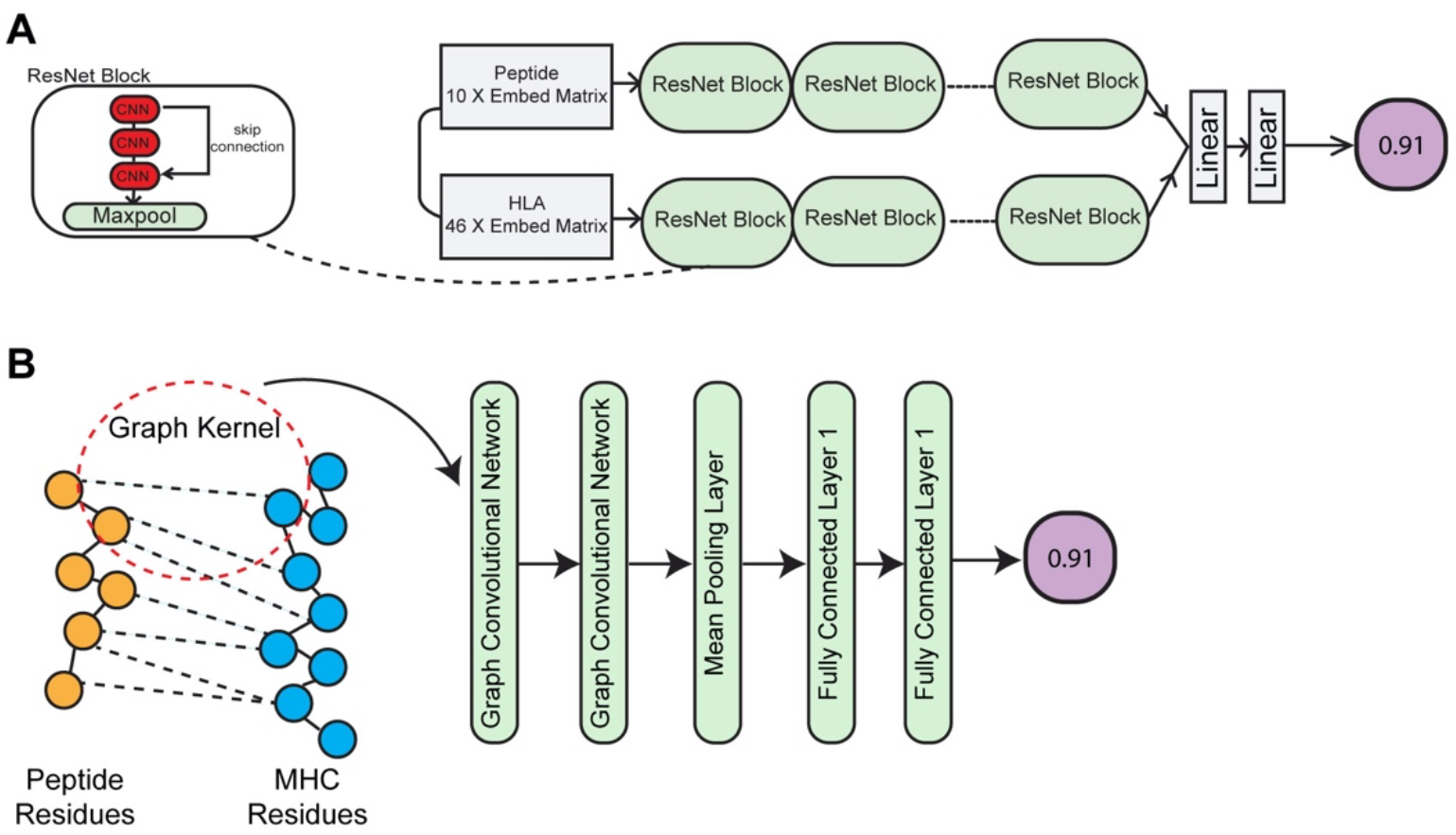
Schematic overview of the ResNet and Graph Neural Network (GNN) models. (A) The ResNet architecture is depicted with three Residual blocks, chained together to extract high-level abstract features associated with immunogenic and non-immunogenic sequences. Each residual block encompasses three convolutional neural network layers followed by a maxpool layer. (B) The GNN architecture is depicted with the first two layers of the graph kernel designed to aggregate neighbors’ attributes (physicochemical properties of each amino acid). A mean pooling layer is used to integrate the graph-level embedding, followed by two dense fully-connected layers to predict immunogenicity.

### Occlusion Sensitivity

To assess the relative importance of each amino-acid position in the model, we sequentially occluded those features associated with each position by setting the values = 0 and re-assessed performance by recording the decrease in resultant predictive score. We measured the performance decrease in all 4,143 positive training instances. We sampled 2,000 positive instances each time and measured the decrease in performance and a rank of position was derived and recorded in an array. Note that we did not retrain the initial model but rather zeroed-out/masked each position. We simulated this process 100 times to validate the robustness of the ranking information. A one-sided Mann Whitney U test was performed to test the statistical significance of each occlusion. The motif heatmap of specific MHC alleles were generated based on the schema proposed by Hu et al. [24], where a position-weighted matrix was produced from all collected immunogenic peptides of the queried MHC allele as described by:

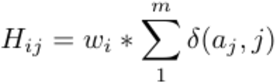

Where **H** is the resultant motif matrix, **w** is the position importance derived from occlusion analysis, **j** denotes 20 amino acids, **i** denotes the position index and **m** signifies the overall number of immunogenic peptides for the queried MHC allele and δ is an indicator function.

### Benchmarking

We benchmarked DeepImmuno-CNN against two existing immunogenicity prediction tools with a high reported auROC. For deep learning based methods, we benchmark against a Gated Recurrent Unit (GRU) [30] based deep learning model DeepHLApan [10]. We downloaded the docker file from docker hub (https://hub.docker.com/r/biopharm/deephlapan) specified in the github page and ran the software in a docker container. We also benchmarked against IEDB’s default MHC-I immunogenicity prediction algorithm [8] from the IEDB web portal. Benchmarking results are shown in **Figure 2**. Other algorithms were excluded for evaluation due to either challenging to use interfaces (e.g. inability to query multiple alleles simultaneously – INeo-Epp [9]) or because they could not be directly compared due to underlying assumptions of the method (e.g., Neopepsee [7]). The evaluated algorithms were not time benchmarked, as the running time for all algorithms were relatively fast (seconds).

**Figure 2.**
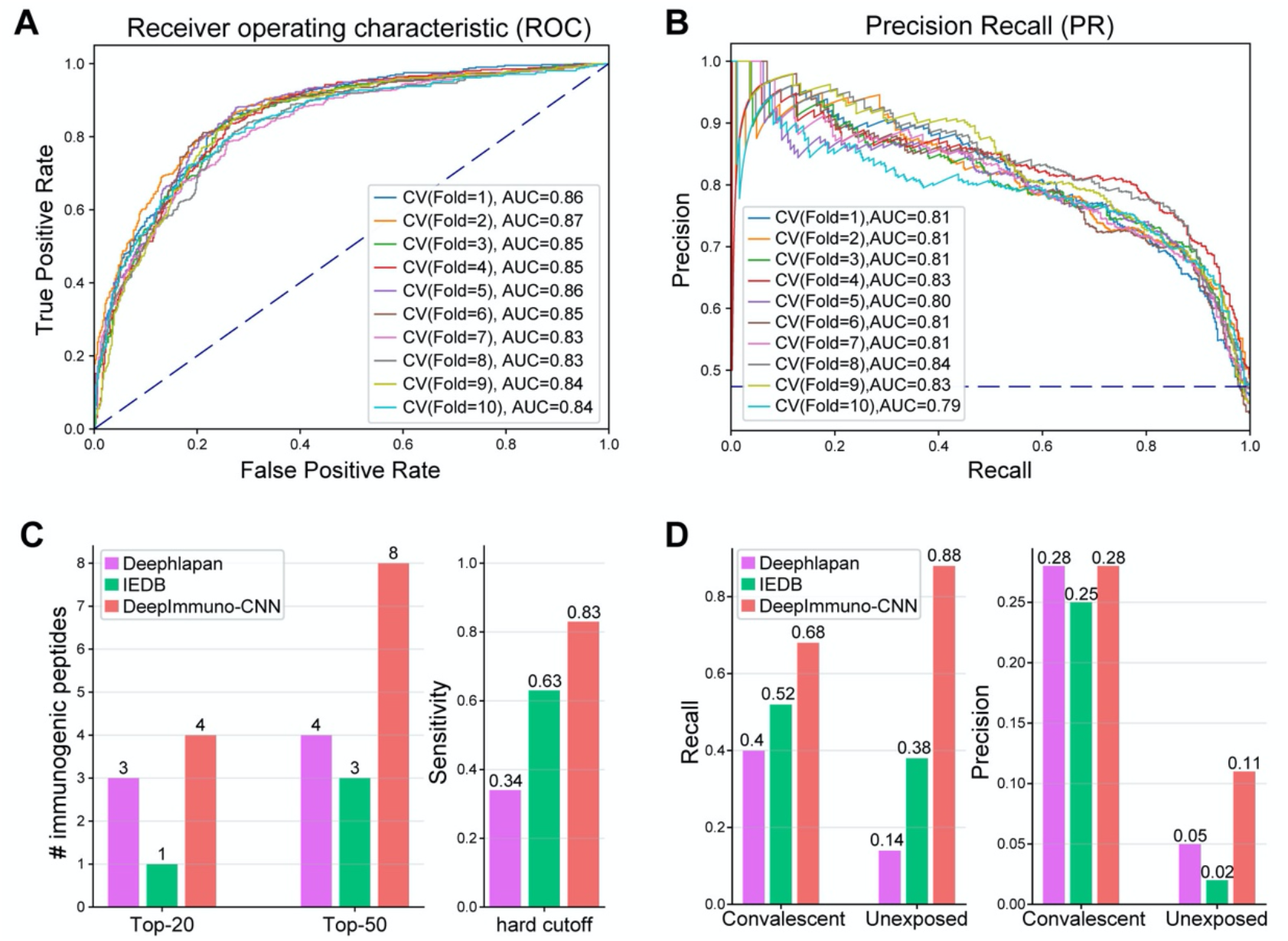
DeepImmuno-CNN produces stable predictions and outperforms existing methods. (A-B) The (A) ROC curve and (B) Precision Recall curve of only DeepImmuno-CNN’s performance on 10-fold validation of the IEDB training dataset. (C) Comparison of immunogenicity predictions from an experimentally validated tumor neoantigen dataset (637 tested), with the number of true positive predictions overlapping with each algorithm’s top 20 or top 50 predictions (left), or the sensitivity of each algorithm using a static scoring threshold (right). (D) In COVID-19 study, recall (left) and precision (right) of each algorithm in convalescent COVID-19 patients and the unexposed individuals.

### Generative Adversarial Network (GAN)

To determine whether immunogenic peptides could not only be predicted but learned and simulated, we trained a GAN model. The GAN model is composed of a generator and a discriminator. We adopted the architecture proposed by Gupta et al [13], as shown in **Figure 1D**. Briefly, an one-hot encoding strategy was used to facilitate the inverse transformation from a probability to pseudo-sequence, then five residual blocks were chained together in both the generator and the discriminator. A 1-dimensional convolutional layer was used to convert the number of channels to be the number of 21 amino acids sequences. We modified the general objective function using Wasserstein distance (WGAN) [17] and improved the stability of training by enforcing 1-Lipshitz constraint using a gradient penalty (WGAN-GP) [31]. The proposed GAN model uses the following loss function:

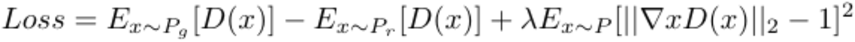

Where **P_g_** is the generated sequence, **Pr** is the real sequence, and **D(x)** indicates the predictive score from the discriminator.

We applied a previously described training strategy for the GAN [13]. Here, gumbel-softmax (tau=0.75) was used in lieu of ordinary softmax to allow sampling from the discrete output. Beta1 and Beta2 hyperparameters of the adaptive learning Adam optimization algorithm were set to 0.5 and 0.9 respectively. Finally, the parameters in the discriminator are updated every mini-batch, while the parameters in the generator are updated every 10 mini-batches. The model was trained using batch_size=64 and trained on 100 epochs.

### Similarity between pseudo-sequence and real sequence

The similarity between two peptides’ sequences was defined as the longest contiguous common sequence length between two queried sequences. For two sequence **S1** and **S2**, the similarity was computed as:

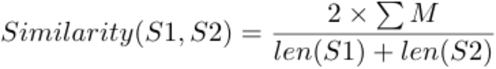

Where **M** denotes the length of each longest common sequence (LCS). **S1** and **S2** belong to 20 amino acids plus a placeholder amino acid “-”. We used the SequenceMatcher function in Python3 difflib package for calculation.

### Web application development

We built an interactive web application (https://deepimmuno.herokuapp.com) for quick query of immunogenic epitopes. The front-end was implemented in HTML5 with bootstrap 4 framework. The back-end was implemented in the Flask python3 framework. The webpage was deployed to Heroku platform through the DeepImmuno github webportal. The weblogos are generated using (http://weblogo.threeplusone.com/create.cgi) for bound peptides of each MHC allele [32]. Please note, if not recently used, the web app takes 30-60 seconds to load for each session.

## RESULTS

DeepImmuno-CNN was developed with the primary objective of improving immunogenicity predictions for relevant disease antigens identified from diverse upstream approaches. To this end, we set out to systematically evaluate existing as well as potential machine and deep learning strategies. This benchmarking was performed on multiple recently described high-quality experimentally validated immunogenic peptides, after carefully excluding low-confidence experimental results (**Methods**).

### Evaluation criteria

We used different evaluation metrics depending on the characteristics of each testing dataset. For the tumor neoantigen test dataset, we considered: a restricted dataset of the (1) top 20 or (2) top 50 immunogenic peptides predictions for each algorithm’s or (3) overall sensitivity. The top 20 or 50 immunogenic peptides were purposely selected as these are the same number of peptides considered in prior related discovery or clinical reports [20]. For the sensitivity analysis, a threshold of 0.5 was used for DeepImmuno-CNN and DeepHLApan and a threshold of 0 for the IEDB default classification algorithm, which has a distinct scoring range. Since an absolute threshold is not used for DeepImmuno-CNN, which outputs a score based on the trained binomial-distribution, this threshold was only used for comparative benchmarking purposes. It is worth noting that we do not consider specificity in the validated neoantigen dataset because each peptide has only been tested in a single cancer patient and hence it is highly likely that a certain peptide can be immunogenic in a larger population with more diverse TCR repertoires.

For antigens from a recent COVID-19 study, we considered recall and precision as the primary criteria due to a much higher number of negative versus positive immunogenic antigens (imbalanced). For evaluation, we used 10-fold cross validation to assess the effectiveness of DeepImmuno-CNN. In each iteration, area under the Receiver Operating Characteristic curve (auROC) and area under the precision recall curve (auPR) were computed to compare performance at different selected cutoffs. auPR is more informative than auROC in an imbalanced scenario due to the incorrect interpretation of specificity [33]. For the five evaluated machine learning algorithms, we tuned the major hyperparameters based on ten-fold cross validation with Root Mean Square Error (RMSE) as the evaluation criterion.

### Comparison of immunogenicity prediction models

To account for variable immunogenic potential for each evaluated peptide, we fitted a beta-binomial probabilistic model in the training dataset to derive a continuous immunogenic score (**Figure 1A, B** and **Methods**). For instance, the peptide RPIDDPFGL for the HLA allele HLA-B*0702 was tested in 40 subjects and triggered a T cell response in all 40 subjects, whereas the peptide KTWGQYWQV in conjunction with HLA-A*0201 elicited a T cell response in only 1 out of 6 subjects, even though both are “immunogenic”. Hence, the former epitope result is of greater confidence. By considering the derived immunogenic potential, we can better ensure that the final predictive scores are more reflective of an epitope’s real immunogenicity.

To select the best predictive model, we constructed five traditional machine learning regressors (ElasticNet, KNN, SVM, Random Forest and AdaBoost) and critical hyperparameters were tuned via cross-validation (**Methods**). In addition, we explored the potential of three deep learning models (CNN, ResNet, GNN). We systematically gauged their performance in three testing datasets (dengue virus[19], tumor neoantigens[20] and SARS-CoV-2 [1]) (**Supplementary Table 2**). Random Forest based regressor had a slightly better RMSE in the nested 10-fold validation than other models, and AdaBoost regression performed the best in dengue virus dataset with average accuracy = 0.91. However, the CNN model achieved superior performance in the neoantigen dataset, where it predicted 2.9 and 5.9 immunogenic epitopes on average and in its top 20 and top 50 predictions, respectively. All the models achieved similar results on the SARS-Cov-2 dataset with an average recall around 0.72 in convalescent patients and 0.81 in the unexposed groups. Given that it is able to mimic the interaction between peptide and MHC, we designed a Graph CNN model, however it suffered from “shortcut learning” [34] such that all the predictive values are around 0.5 to achieve a lower loss during the training stage. This can be attributed to the fact that the explicit weight assignment in the graph may not entirely reflect the real peptide-MHC interactions, which in turn can lead to ambiguous results. To explore whether increasing the complexity of the neural network architecture can boost performance, we constructed a ResNet model, with 12 layers and skip connections. As ResNet did not increase the performance and had inferior results in 8 out of 9 evaluation criteria across three testing datasets, we surmise that a more complex model is not required. Considering its performance overall and in human disease datasets, adaptability to training datasets of variable size and the complexity of the model, we chose CNN as the optimal prediction model for further analysis, which we call hereafter DeepImmuno-CNN. As a final consideration, we attempted to validate our proposed amino acid encoding strategy which considers both indices derived from amino acid physicochemical properties (AAindex) and HLA allotype information (paratopes). While, use of these algorithms did not result in significant performance boosts with neural network based approaches over alternative strategies, our selected encoding methods did not decrease performance and did offer a performance boost for specific machine learning methods (Random Forest) for specific test datasets, suggesting its benefits may be situation dependent (**Supplementary Table 2**, **Supplementary Figure 3**).

**Supplementary Figure 3.**
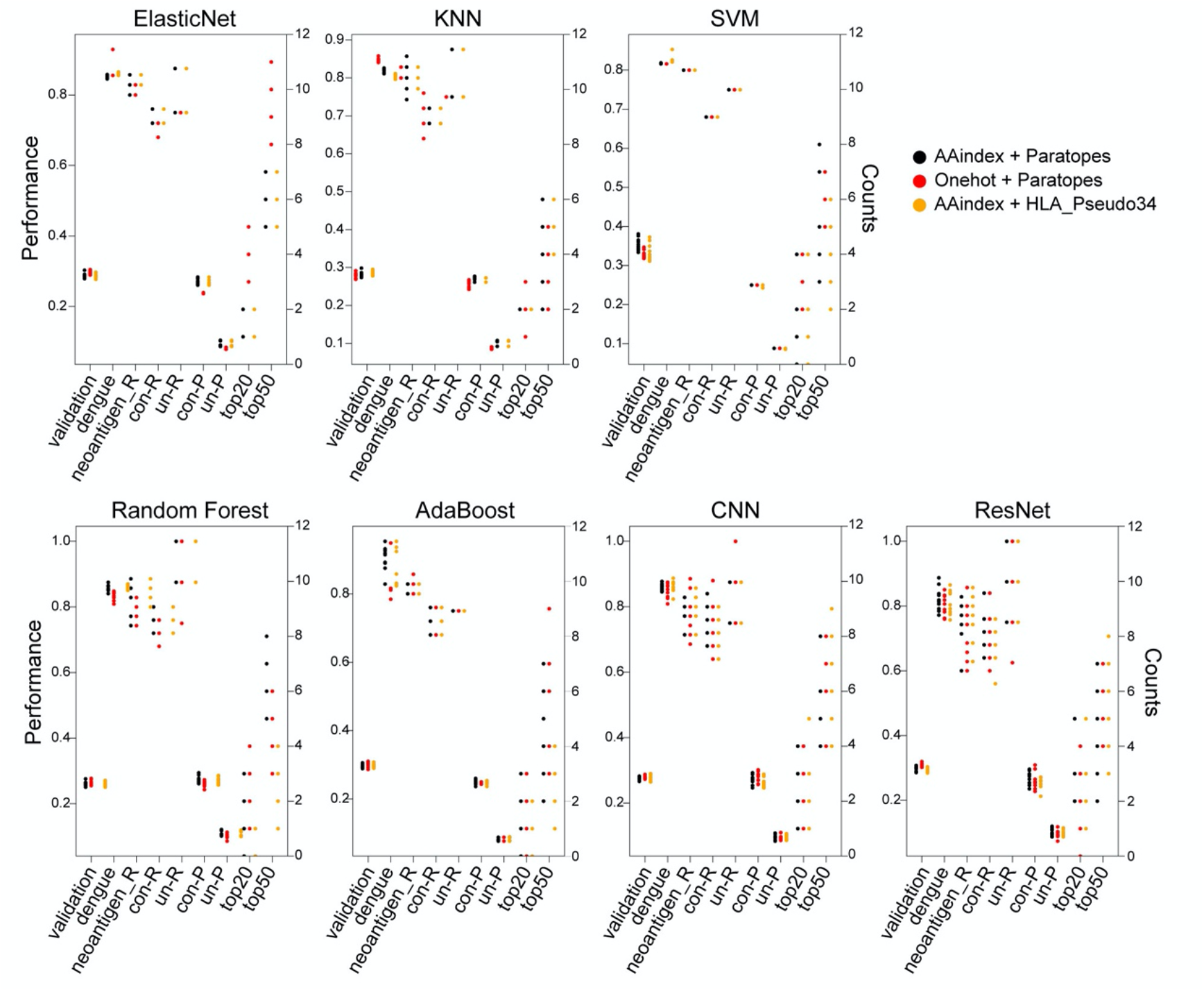
Immunogenicity prediction following an ablation test of different encoding strategies across 7 evaluated algorithms. Relative immunogenicity detection performance of seven evaluated algorithms (ElasticNet, KNN, SVM, Random Forest, AdaBoost, CNN and ResNet) and three different encoding strategies (AAindex+Paratopes, One-hot encoding + Parartopes and AAindex + HLA Pseudo34). AAindex encoding and HLA paratopes representation is shown in black, Onehot encoding and HLA paratopes representation is shown in red, AAindex encoding and HLA pseudo34 sequences representation is shown in orange. The x-axis represents nine different performance evaluation statistics across the four test datasets. These statistical metrics are: 1) validation (RMSE in nested 10-fold validation dataset), 2) dengue (Accuracy in dengue virus dataset), 3) neoantigen_R (Recall in cancer neoantigen dataset) and 4) con-R (Recall in COVID-19 convalescent patients group), un-R (Recall in COVID-19 unexposed patients group), con-P (Precision in COVID-19 convalescent patients group) and un-P (Precision in COVID-19 unexposed patients group). top 20 = immunogenic neoantigen in top 20 ranked hits; top 50 = immunogenic neoantigen in top 50 ranked hits. The indicated dataset-specific performance metric is indicated on the y-axis (range 0-1, left) or top 20/50 hits (counts, right).

To validate the effectiveness of the DeepImmuno-CNN model, we conducted a ten-fold cross validation in the IEDB dataset, on its own (**Figure 2A, B**). We foundDeepImmuno-CNN to be highly stable with a high average auROC (0.85) and auPR (0.81) for each fold. We next compared the performance of this CNN model relative to other prior described immunogenicity prediction methods, specifically DeepHLApan and IEDB (default algorithm), as these methods are well-validated and have easy-to-use interfaces. When evaluated in the tumor neoantigen dataset, DeepImmuno-CNN found an impressive 29 out of 35 (83%) immunogenic neoantigens, relative to IEDB which found 63% and DeepHLApan which only found (34%) out of a total of 637 antigens experimentally tested (**Figure 2C**). For the same neoantigen dataset, DeepImmuno-CNN predicts 4 in the top 20 and 8 in the top 50 neoantigens, while IEDB performed relatively poorly (1 in the top 20 and 4 in the top 50), with DeepHLApan producing intermediate results (**Figure 2C**).

We further evaluated DeepImmuno-CNN using a recently published COVID-19 study, where immunogenic peptides were validated from two groups of subjects. Convalescent patients have already been infected by SARS-Cov-2 and are in the process of recovering, while unexposed patients haven’t contracted the disease. In both convalescent and unexposed groups, DeepImmuno-CNN achieved the highest sensitivity (68% in convalescent, 88% in unexposed) compared to IEDB (52% in convalescent, 38% in unexposed) and DeepHLApan (40% in convalescent, 14% in unexposed) (**Figure 2D**). DeepImmuno-CNN also achieved the highest precision (0.28 in convalescent, 0.11 in unexposed), with an overall low precision due partially to the fact that COVID-19 patients are a highly selective group and their unique immune profile might not be representative of the whole population. We next looked for potential immunodominant regions in the SARS-Cov-2 proteome, which can be exploited for T cell vaccine development. While our result suggests that both 9-mers and 10-mers do not predict immunodominant regions in general (**Supplementary Figure 4**), some peptides derived from ORF2 spike protein display high immunogenic potential (mean>0.75). These peptides likely reflect the protein’s primary function, which is to interact with human ACE2 receptor [35] and increase the likelihood of triggering a T cell response.

**Supplementary Figure 4.**
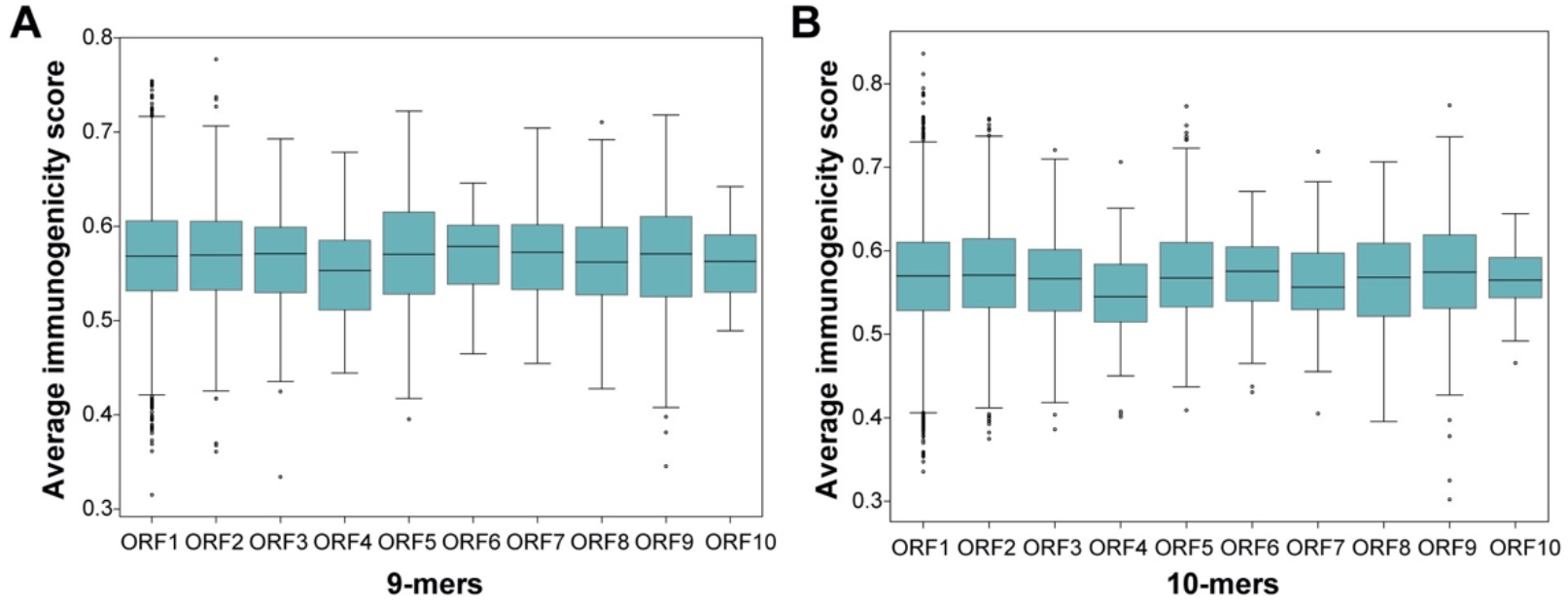
Predicted immunogenicity is consistent across all ORFs in the SARS-Cov-2 proteome. (A) Predicted immunogenicity of all 9-mer peptides translated from 10 different ORFs. (B) Predicted immunogenicity of all 10-mer peptides translated from 10 different ORFs.

### DeepImmuno-CNN reveals salient positions interacting with the TCR

To understand the molecular underpinnings of DeepImmuno-CNN we examined the dependency of this model on each residue position using occlusion sensitivity. The largest decrease in performance corresponds to the most important position across the peptide as shown in a saliency heatmap (**Figure 3A**). We simulated this process 100 times and an ascending ranking was performed each time to highlight the most salient position, as shown in (**Figure 3B**). This analysis reveals that amino acid positions P4 (residue 4), P5 and P6 are consistently the most dependent positions, followed by P2, P8 and P9. Occlusion of the first and second most dependent positions (P4 and P5) compared to the least (P3 and P1) resulted in a significant performance drop of each single positive instance (One-sided Mann-Whitney U test, P-value = 7.9e-209), further evidencing these predictions (**Figure 3C**). These studies support prior structural prediction studies which show that P4-6 interact with the TCR with greatest frequency [36,37], whereas, P2 and P9 serve as anchor points for binding of the peptide-MHC complex [8] and mirrors other computational predictions [5][8].

**Figure 3.**
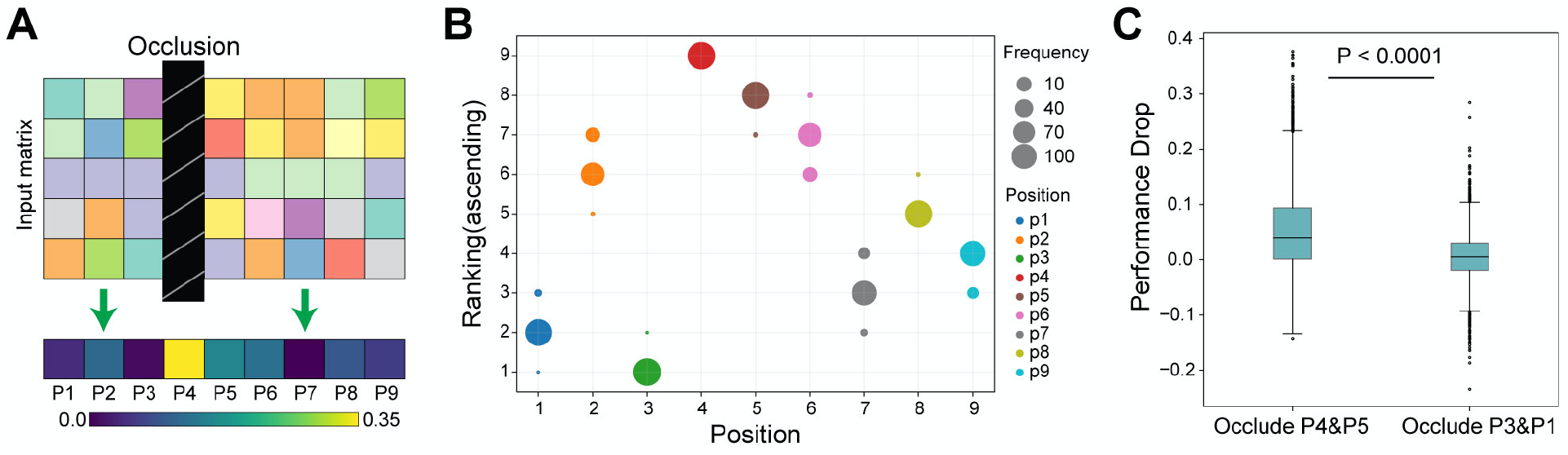
Identification of salinent immunogenic features of peptide-TCR interactions. (A) Schematic overview of the occlusion sensitivity technique to determine the relative contribution of each antigen residue for the DeepImmuno-CNN model predictive score. (B) Ascending importance-rank of each position, with the position with the largest performance drop received the highest ranking across 100 simulations. Dot size corresponds to the frequencies of each position being assigned the denoted rank, with different colors indicating different amino acid positions. (C) Performance drop for the occlusion of P4 + P5 with occlusion of P3 + P1. Onesided Mann-Whitney U test p-value (p=7.94e-209).

To assess the rules governing T cell immunogenicity for different HLA alleles, we next evaluated MHC allele dependence on specific amino acid preferences. To perform this analysis, we collected all immunogenic peptides bound with each allele and derived a motif matrix based on the inferred position importance weight in the model (**Methods**). These results are summarized in **Supplementary Figure 5**. For example, when examining the allele HLA-A*0201, we find Leucine is the most abundant amino acid in position 2 from the model, which is consistent with prior structural evidence [38]. Similarly, in a previous study by Hu et al [24], positions 2 and 9 were predicted to act as anchor points for interactions with this specific HLA allele. Here, our motif matrix additionally suggests that position 4 and 5 interact with the TCR on the other side. We conducted the same analysis on three other HLA alleles (HLA-A*2402, HLA-B*0702, HLA-B*0801). These alleles were chosen because the number of associated immunogenic peptides bound to these three alleles are greater than 150, suggesting that the immunogenic motif matrix for these alleles is stable. As expected, position 4 also shows a stronger pattern across these three alleles, compared to other positions, supporting a similar model of HLA-TCR interactions.

**Supplementary Figure 5.**
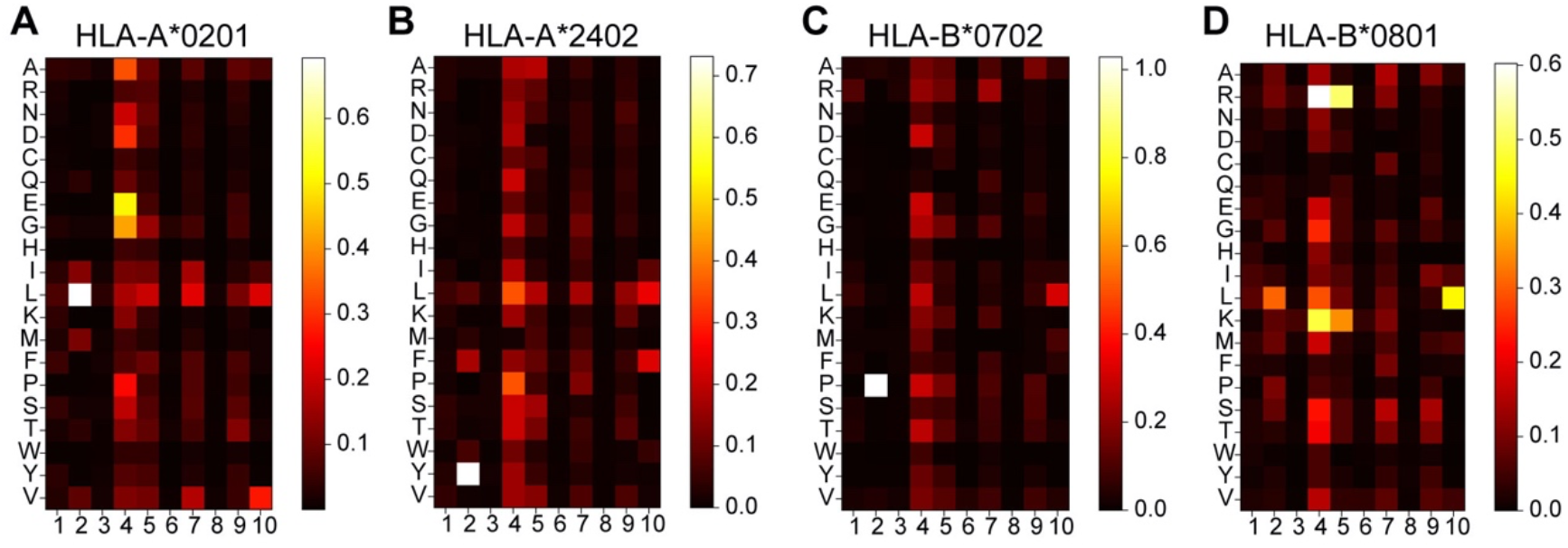
Immunogenicity motif analysis reveals critical HLA-peptide-TCR interacting sequences. The immunogenic motif heatmap for (A) 2,046 HLA-A*0201, (B) 210 HLA-A*2402, (C) 162 HLA-B*0702 and (D) 176 HLA-B*0801 bound epitopes in the IEDB training dataset. Positions are weighted by their relative importance derived from the occlusion sensitivity analysis in Figure 3.

### Deeplmmuno-GAN accurately mimics immunogenic peptide sequences

To better understand the molecular interactions and biochemical properties of T cell immunogenicity, we attempted to generate *de novo* immunogenic peptides using a GAN-based approach. Successful creation of such peptides would indicate that immunogenic sequence motifs are learnable, potentially paving the way for direct synthesis and optimization of peptides for diverse applications (e.g., enhanced immunogenicity)[39].

As a proof-of-concept, we collected all immunogenic peptides known to bind to HLA-A*0201 (the most abundant allele in the training database) for training the deep GAN model. We trained a Wasserstein GAN model for 100 epochs (**Methods**) and extracted the generative pseudo-sequences from every 20 epochs. We utilized the same encoding schema we used in the prediction model to perform dimension reduction using PCA and visualized the distribution of generative and real immunogenic sequences **(Figure 4A,B** and **Supplementary Figure 6A**). When viewed as a PCA projection, we find that random peptide sequences significantly deviate from the experimentally validated immunogenic peptide sequences, prior to GAN model training. However, after GAN model training, the generative pseudo-sequence maps to a common coordinate embedding within the PCA projection to that of real immunogenic peptide sequences. These data suggest that the GAN model is able to extract the high-level features from real instances and teach the generator to output similar immunogenic peptides built from random sequence as a starting input. The same distribution shifts were observed with tSNE dimensionality reduction (**Supplementary Figure 6B**).

**Figure 4.**
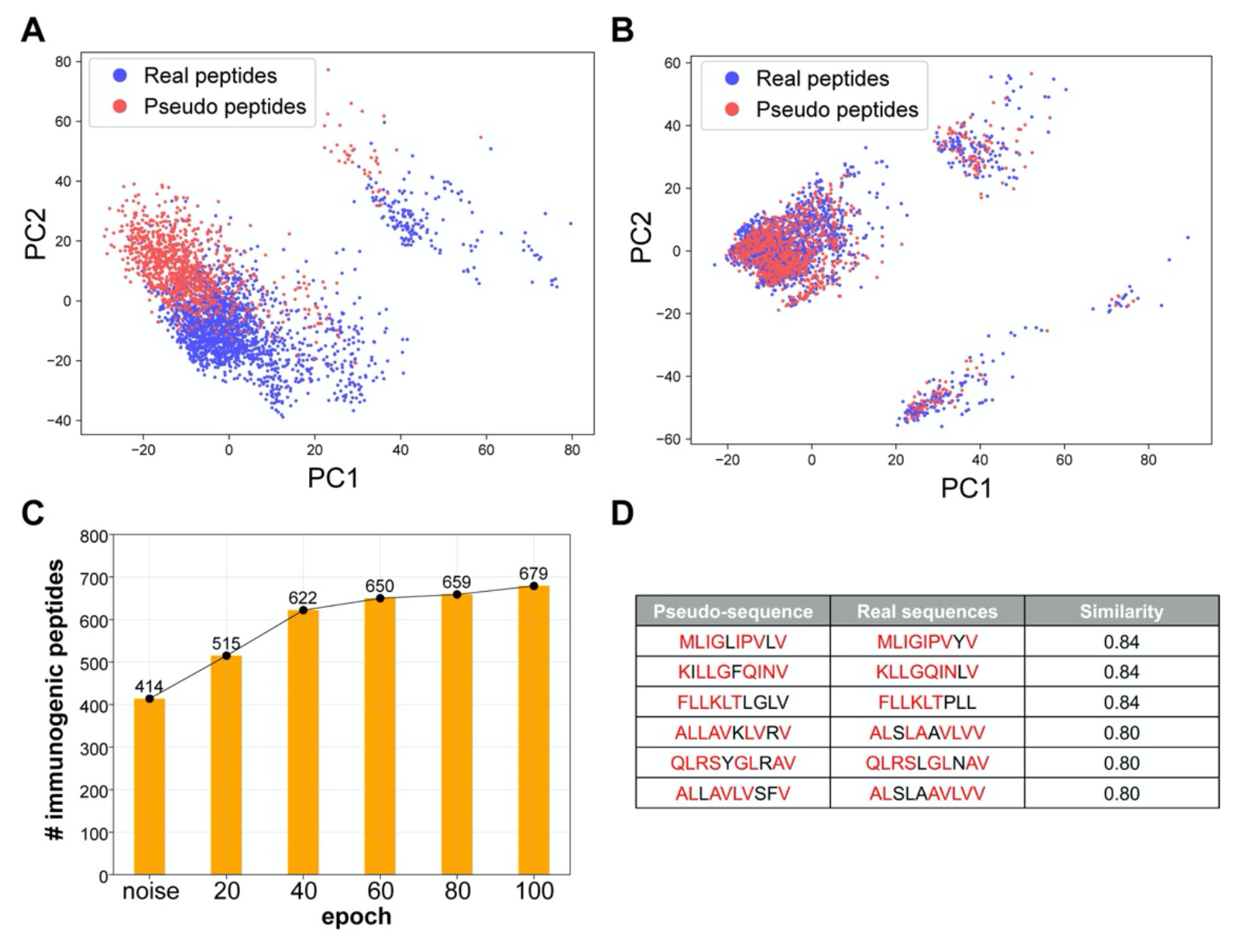
DeepImmuno-GAN is able to learn and produce synthetic immunogenic pseudo-sequences. (A-B) PCA analysis of the distribution of real sequences (blue dots) and random generative sequences (red dots) (A) prior to training and (B) after training (100 epochs). The degree of common embedding is considered an indicator of prediction similarity. (C) The number of DeepImmuno-CNN predicted immunogenic peptides, produced from noise, in different GAN training epochs. (D) Example generative pseudo-sequences and their most similar counterparts in experimentally observed HLA-A*0201 immunogenic peptides.

To further assess the immunogenicity of these generative sequences, we submitted all generated sequences at different epoch points to our DeepImmuno-CNN model. At the beginning, the 1,024 random sequences were found to only contain 40% of the immunogenic sequence (predictive score > 0.5). As training progresses, the fraction of immunogenic peptides gradually increases to 67%, which translates to 265 more immunogenic peptides generated during training (**Figure 4C**). We compared each generative pseudo-sequence to their most similar real counterparts (**Figure 4D**). The similarity was defined as the total longest contiguous matching subsequence (LCS) between the real and pseudo-sequence, with 87% (891/1024) of all pseudo-sequences having >60% similarity to their matched real immunogenic peptides (**Methods**) (**Supplementary Figure 7**) [40]. Hence, immunogenic peptides can be learned and produced when sufficient training data exists.

**Supplementary Figure 6.**
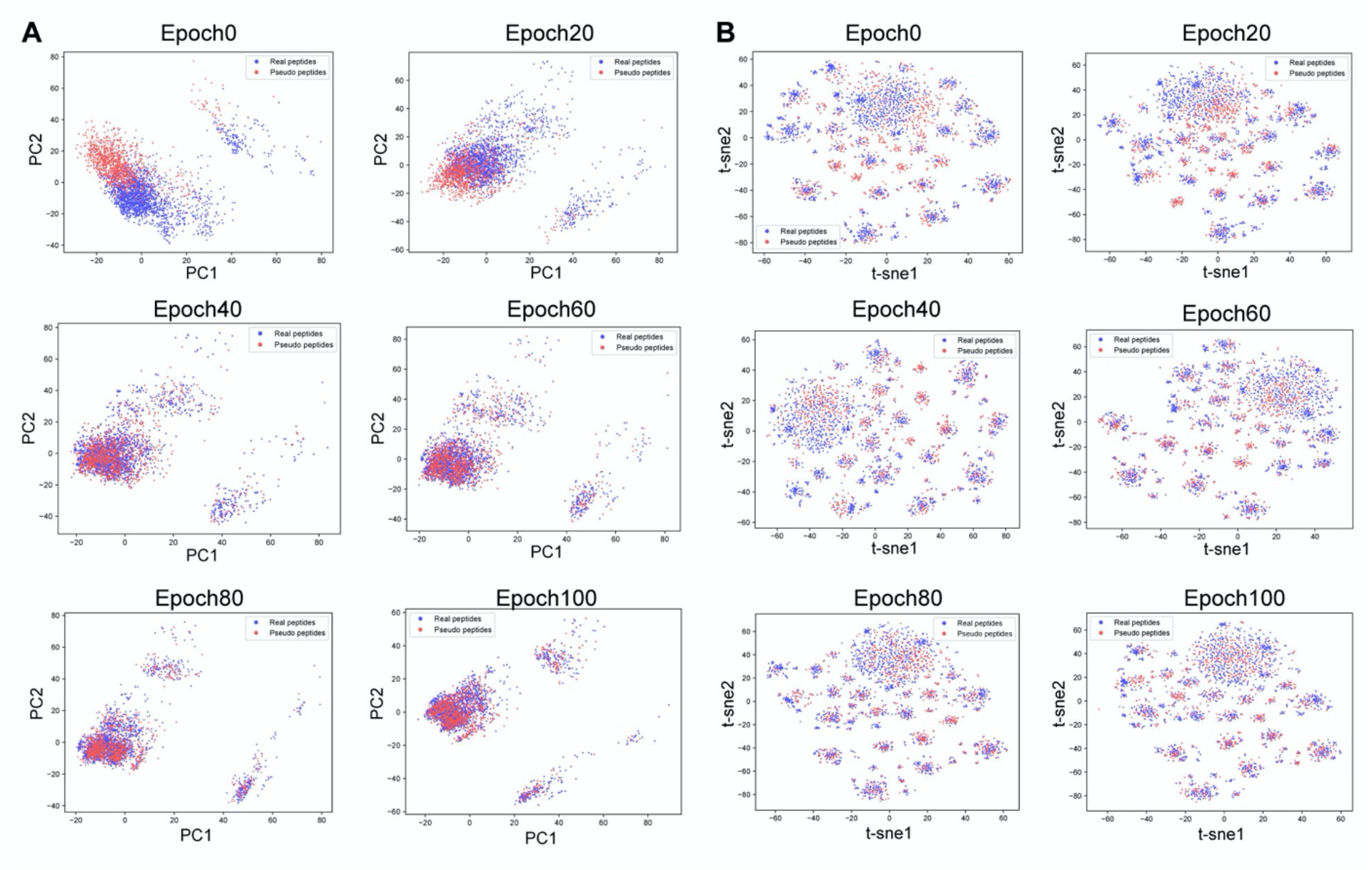
Convergence of real immunogenic and pseudo-sequences with progressive GAN training. GAN generative sequences from epoch 0, epoch 20, epoch 40, epoch 60, epoch 100 were concatenated with real HLA-A*0201 instances and their joint embedding spaces were visualized using either (A) PCA or (B) t-SNE.

**Supplementary Figure 7.**
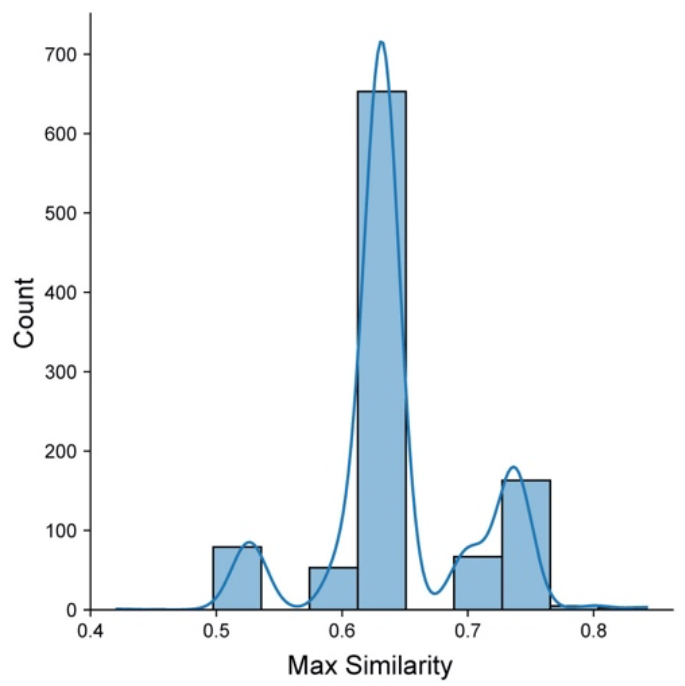
Distribution of max similarity between GAN generated pseudo-sequences and their matched real immunogenic peptides in HLA-A*0201. The maximum similarity for each GAN-generated pseudo-sequence and its most similar counterpart in real immunogenic peptide repertoires are shown, with similarity defined as the longest contiguous common sequence length in total between two queried sequences (**Methods**).

### Online web interface

In order to simplify the process of building DeepImmuno from source code, we developed an easy-to-use web interface allowing users to quickly query peptide sequences to predict immunogenicity potential for a given HLA allele. Additionally, this service allows a user to query for which HLA allele would yield the highest immunogenicity and hence which patients might benefit most from an immunogenic therapy. A third supported query type is for an HLA allele, what epitopes it will prefer or disfavor. To address the latter question, the user can simply enter the queried epitope sequence and HLA allele to obtain the immunogenicity score, top five combinations with different HLA alleles and a weblogo view of all immunogenic and non-immunogenic epitopes associated with a certain HLA allele (**Figure 5**). Moreover, the DeepImmuno web portal allows users to perform multiple queries by specifying an input file with epitope sequence information and an output text file with the predicted immunogenicity scores will automatically be returned.

**Figure 5.**
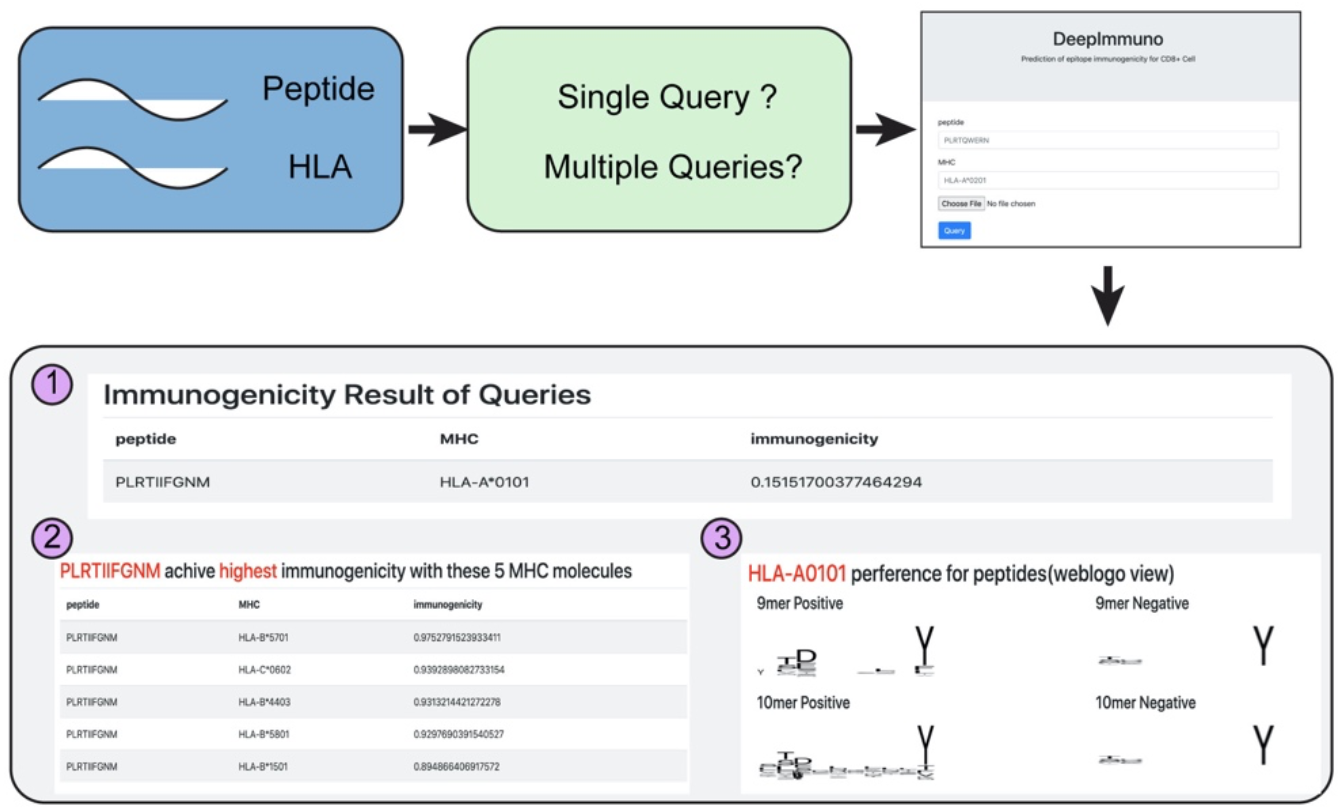
The DeepImmuno web interface. An easy-to-use web interface for querying peptide and HLA sequence pairs. The three primary outputs of the interface are: (1) Immunogenicity score for queried peptide-HLA combination, (2) the top 5 HLA combinations that will yield the highest immunogenicity score for each queried peptide and the (3) preferential motif of the queried HLA allele. Please note, if not recently used, the web app takes 30-60 seconds to load for each session.

## DISCUSSION

The accurate identification of potential immunogenic epitopes remains a significant challenge for understanding the molecular mechanisms underlying host immune response and designing effective targeted therapies. Given the fact that millions of possible epitopes can be generated from human protein coding genes [3], experimentally validating all possibilities is simply not yet feasible. Effective computational models can largely accelerate this process by providing a pre-screening platform to find high-confidence immunogenic epitopes or to eliminate low confidence predictions. Machine learning and deep learning algorithms have been shown to provide increased performance in a wide spectrum of bioinformatics applications [41,42]. However, comprehensive benchmarking and the selection of an optimal encoding strategy are required to develop improved models that can be applied to diverse testing datasets.

In this manuscript, we developed a beta-binomial model to generate more accurate immunogenicity potential by considering the overall quality of each experimentally tested antigen in the training dataset. Using these optimized training datasets, we systematically bencharked well-established machine learning and deep learning, and encoding strategies on independent immunogenic disease datasets, to understand the different situations in which these methods boost, decrease or do not impact overall classification performance. From this extensive comparison analysis we found that a CNN model in combination with a physiometric-aware encoding strategy balanced performance across diverse test datasets, while staying robust for different training dataset sizes. Indeed, we found that increasingly complex deep learning models, such as ResNet, could result in overfitting in this specific application. Our DeepImmuno-CNN model was able to significantly outperform two existing highly used immunogenicity prediction workflows, in terms of overall sensitivity and the top ranked hits, when applied to diverse real-world immunogenic antigen datasets, including cancer and COVID-19 infection. From a neoantigen pre-screening perspective, DeepImmuno-CNN, is mostly likely to increase the sensitivity for detection of valid neoantigens, such as tumor-specific mutations or splicing neojunctions, from large-scale genomics assays to be tested in downstream assays. Using this optimized model, we were able to effectively identify the most salient residues for interactions between peptide-MHC and TCR, which were recapitulated and added to prior knowledge. Moreover, we developed a GAN modelling approach to accurately generate immunogenic peptides from random noise and demonstrated the biochemical interactions were learnable given sufficient training data.

Despite these advances described herein, several challenges remain in the field of immunogenicity prediction. While our model significantly improves upon existing approaches in terms of sensitivity, precision and recall, it is noteworthy that all existing approaches remain challenged by lower than preferred specificity to select immunogenic antigens with high confidence. This limitation could be due to the fact that few disease antigens have been thoroughly tested for their ability to mount a T cell response in large patient cohorts to ensure reproducibility and HLA allele coverage. However, it is noteworthy that an indispensable component of epitope recognition is the sequence of the TCR, which has not been taken into consideration due to the fact that there exists few matched TCR sequencing data for forming a sufficiently powered training set [43,44]. In addition, a model incorporating TCR information is only applicable following sufficient deep TCR repertoire patient sequencing. Although new high throughput methods for single-cell TCR sequencing have been developed, such techniques are still infrequently performed in research and clinical settings. The increased use of such techniques are likely to aid in the development of more accurate predictive models. In addition, neoantigen T-cell responses can significantly vary from patient-to-patient, due to a variety of factors including immune cell repertoire differences that impact diversity of activated T cell clones [45,46,47]. Hence, validated immunogenic epitopes may be ineffective in a subset of patients. The ambiguity of the definition of immunogenicity can account for part of the false positive predictions which might in fact be immunogenic for a set of patients. Integrating patients’ immune profiles information and identifying how active the host immune system is can be a valuable extension to current immunogenicity models. Beyond providing a rubric for the design of peptide-related models, we believe our approach can be significantly extended to encode additional variables, such as TCR sequence heterogeneity and can be generalized to address diverse sequence-predictive analyses, beyond immunogenicity.

## Conflict of interests

The authors report no significant conflicts of interest.

## Funding

This work was supported by the Cincinnati Children’s Hospital Research Foundation and funding from the National Institutes of Health [R01CA226802].

## Notes

https://github.com/frankligy/DeepImmuno

### Competing Interest Statement

The authors have declared no competing interest.

